# High-affinity free ubiquitin sensors as quantitative probes of ubiquitin homeostasis and deubiquitination

**DOI:** 10.1101/528711

**Authors:** Yun-Seok Choi, Sarah A. Bollinger, Luisa F. Prada, Francesco Scavone, Tingting Yao, Robert E. Cohen

## Abstract

Ubiquitin (Ub) conjugation is an essential post-translational modification that affects nearly all proteins in eukaryotes. The functions and mechanisms of ubiquitination are areas of extensive and ongoing study, and yet the dynamics and regulation of even free (i.e., unconjugated) Ub are poorly understood. A major impediment has been the lack of simple and robust techniques to quantify Ub levels in cells and to monitor Ub release from conjugates. Here we describe the development of avidity-based fluorescent sensors that address this need. The sensors bind specifically to free Ub, have K_d_ values down to 60 pM, and, in concert with a newly developed workflow, allow us to distinguish and quantify the pools of free, protein-conjugated, and thioesterified forms of Ub from cell lysates. Alternatively, free Ub in fixed cells can be visualized microscopically by staining with a sensor. Real-time assays using the sensors afford unprecedented flexibility and precision to measure deubiquitination of virtually any (poly)Ub conjugate.

## Introduction

In ubiquitination, free Ub is activated by formation of a C-terminal thioester first with an E1 Ub-activating enzyme and then an E2 Ub-conjugating enzyme before it is transferred to substrates, usually to form an isopeptide bond with a protein lysine ε-amine^1^. Deubiquitinating enzymes (DUBs) can disassemble Ub–protein conjugates and recycle Ub; thus, cells contain three classes of Ub: free, thioester-activated, and (iso)peptide conjugated. Because ubiquitination contributes to the regulation of nearly every cellular process, the availability of free Ub needs to be tightly controlled to maintain cell fitness. Mechanisms contributing to Ub homeostasis include its expression as Ub–protein fusions with ribosome subunits and as polyubiquitin, processing of these precursors to generate free monoUb, and finally recycling from Ub–protein conjugates by DUBs^2^. Perturbations of these processes can deplete cellular free Ub and cause defects in cell development or neuronal functions^2–9^and inhibit proliferation in several cancer cell lines^10,11^. Conversely, transgenic mice that overexpress Ub by just 2 or 3-fold exhibit neurological abnormalities^12^.

Although the need to maintain and regulate intracellular free Ub is now well established, studies of Ub levels have been hampered by the lack of reagents for specific and quantitative measurements of distinct Ub pools. One approach to monitor intracellular Ub has been ectopic expression of GFP-tagged versions of Ub^13^, but interpretations of results from such experiments can be compromised by perturbations to the regulation of endogenous Ub and non-physiological behavior of the tagged Ub. Typically, to quantify endogenous free, conjugated, or total Ub, anti-Ub antibodies in conjunction with ELISA or SDS-PAGE and immunoblotting are used. However, with respect to sensitivity and reliability of quantitation, those approaches have major drawbacks. For example, due to the extremely high structural diversity of polyUb and Ub–protein conjugates^14^, even with monoclonal anti-Ub antibodies, binding efficiencies will vary among the many different forms of Ub in cell lysates. Moreover, particularly with quantitation by western blots, dynamic range is inherently very limited and reproducibility can be difficult to achieve. More recently, Ub Protein Standard Absolute Quantification (Ub-PSAQ) mass spectrometry (MS) has been described to quantify free and conjugated Ub from cell lysates^15^. However, this method does not resolve the pool of thioester-activated Ub, and its dependence on sophisticated instrumentation and sequential affinity-based isolation steps makes it challenging to implement for most laboratories. Finally, none of the aforementioned approaches to Ub quantitation are amenable to real-time measurements of free Ub concentrations as they change, for example, during enzyme-catalyzed deubiquitination reactions.

With the dual goals of having a simple, reliable method to quantify cellular Ub pools and a sensitive and versatile real-time DUB assay, we embarked on development of sensors for free Ub detection and quantitation. To distinguish the free, activated, and conjugated Ub pools, we developed protocols to convert either thioesterified or (iso)peptide-conjugated Ub into free Ub for subsequent quantitation with a sensor reagent. To engineer the sensors, our general strategy was to fuse genetically two or three Ub binding domains (UBDs) of known structure that bind to non-overlapping Ub surfaces, and to exploit avidity effects to achieve high affinity and selectivity. To convert the binding proteins into sensors, we attached fluorescent dyes whose intensity changed in response to binding by free Ub. We demonstrate that the sensors provide convenient means to (i) measure free, activated, and conjugated intracellular Ub; (ii) quantify deubiquitination of unlabeled (poly)Ub–protein conjugates in real-time DUB assays; and (iii) localize and quantify endogenous free Ub by fluorescence microscopy of fixed cells.

## Results

### Design and characterization of the sensors

The free Ub sensors were developed to have high affinity and selectivity for free Ub and report binding events via a strategically placed fluorophore. Our design strategy was to assemble Ub binding proteins from multiple UBDs linked in tandem; peptide linkers were kept to a minimum length in order to promote avidity while minimizing the entropic cost of binding. Most UBDs bind Ub through interactions with one of three different surfaces: the Ub hydrophobic patch surrounding residue I44, the Ub C-terminal tail, and the surface around D58 (Fig. 1a, upper panel). Individually, UBDs bind Ub with only modest affinity (K_d_ = 10^−5^ to 10^−3^ M), but by linking two or three weak-binding UBDs that target distinct Ub surfaces, we expected that high affinity could be achieved overall. An early version of such an avidity-based binder that we call tIVR employed IsoT^Buz^, Vps27^UIM^, and Rabex5^Ruz^ domains fused in tandem with flexible peptide linkers (Fig. 1a, lower left panel and Supplementary Fig. S1). The IsoT^Buz^ domain (K_d_ = 3 μM for Ub)^16^, which binds primarily to residues at the Ub C-terminus, conferred selectivity for free Ub, whereas the Vps27^UIM^ (K_d_ = 117 μM)^17^ and Rabex5^Ruz^ (K_d_ = 12 μM)^18,19^ domains worked synergistically with IsoT^Buz^ to increase overall affinity and specificity for Ub.

**Fig. 1.**
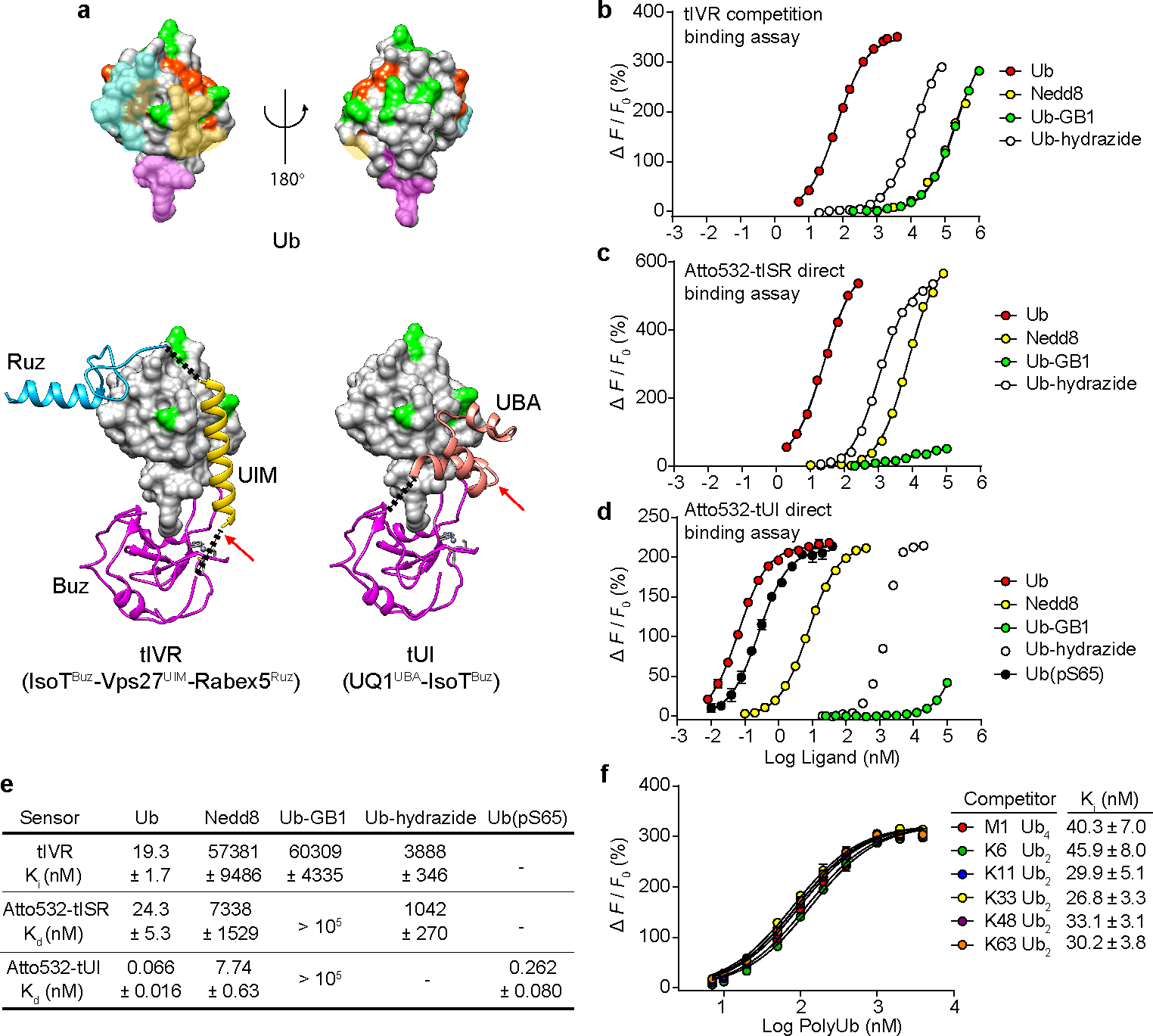
Sensor design and characterization. **a**, Ub (upper panel) has distinct surfaces recognized by three classes of UBDs. Ub and UBDs are shown in surface and ribbon representations, respectively. For Ub (upper models), surfaces where Buz, UIM or UBA, and Ruz domains bind are in magenta, yellow, and cyan, respectively. Lysine and M1 sidechains (green) and phosphorylation sites (orange) are highlighted. Ub complexes with tIVR (lower left) and tUI (lower right) were modeled using individual UBD-Ub complex structures (PDB 2G45, 2FIF, 1Q0W, and 2JY6). The black dotted lines indicate linkers installed to connect UBDs, and red arrows show sites of fluorophore attachment. **b**, tIVR affinities for Ub and UbL derivatives were measured by competition with 1 nM Atto532-Ub(S20C) in the presence of 6 nM tIVR. ∆*F* is the fluorescence intensity change of Atto532-Ub(S20C) upon addition of competitor and *F*_0_ is the fluorescence without competitor. Fluorescence intensity changes of **(c)** Atto532-tISR or **(d)** Atto532-tUI were measured by direct titrations with the Ub or UbL derivatives indicated and fit with a 1:1 binding model as described in Methods. **e**,Affinities (K_d_ or K_i_) of the three sensors determined for the indicated Ub and UbL ligands. **f**, Effects of Ub–Ub linkage type were assessed from competition binding assays with 0.8 nM Atto532-Ub(S20C) and 6.0 nM tIVR titrated with 7 to 4000 nM of the indicated polyUb ligands. Statistical errors listed are standard deviations from the fits. Note that in **d**, the titration with Ub-hydrazide shows a saturation binding curve that might be due to trace contamination by free Ub; therefore, a K_d_ value is not shown. Small error bars in **d** and **f**, determined from duplicate assays, are masked by the point symbols.

To measure the affinity between tIVR and free Ub, we first titrated a fluorescent Ub derivative (Atto532-Ub(S20C); see Methods) with tIVR and determined a 0.4 nM K_d_ based on an observed 3.5-fold fluorescence decrease upon formation of the 1:1 complex (Supplementary Fig. S2a). Then, by taking advantage of the high affinity and fluorescence intensity change of Atto532-Ub(S20C) upon binding to tIVR, we determined tIVR affinities for Ub and other ligands by use of competition binding assays (Fig. 1b). We found that free Ub binds to tIVR with a K_d_ of 19 nM (measured as a K_i_; Fig. 1b); the nearly 50-fold stronger binding of Atto532-Ub(S20C) indicates that the Atto532 dye contributes favorably (ΔG ~ −0.2 kcal/mol) to the interaction with tIVR. The competition assays additionally showed that tIVR has high selectivity against the Ub-like (UbL) protein Nedd8 and against Ub derivatives that lack a free C-terminal carboxylate (Fig. 1b). Among the many different UbL proteins found in eukaryotes^20^, Nedd8 is most similar to Ub in its sequence and tertiary structure, and its C-terminal four amino acids are identical^21^; nonetheless, tIVR has a 3000-fold preference for Ub. A similarly large discrimination was observed against Ub-GB1, a mimic of a Ub–protein conjugate in which the Ub C-terminus is extended by the Protein G 56-amino acid B1 domain, and even addition of the small adduct hydrazine to the Ub C-terminus decreased tIVR affinity 150-fold relative to free Ub (Fig. 1b).

In order to provide the sensor with a direct readout of Ub binding, we explored site-specific labeling with fluorescent dyes (Supplementary Fig. S3). We found that tIVR modified with Atto532-maleimide on C130 showed a 3-fold fluorescence increase upon Ub binding (Supplementary Fig. S2b). By replacing the Vps27^UIM^ with the S5a^UIM^ domain and introducing two amino acid substitutions in the IsoT^Buz^ domain (Fig. 1a and Supplementary Fig. S1), we developed a second-generation sensor, tISR, that binds to free Ub nearly 10-times tighter than tIVR (Supplementary Fig. S2c,d). Atto532-labeled tISR showed a 6-fold fluorescence increase upon binding of free Ub, but the fluorophore reduced affinity for Ub so that Atto532-labeled tISR and unlabeled tIVR showed similar K_d_ values of 24.3 nM and 25.4 nM, respectively (Fig. 1c,e and Supplementary Fig. S2b).

A third generation of free Ub sensor was developed with the goal of achieving even greater affinity. Ubiquilin-1 UBA (UQ1^UBA^) was used to replace the tIVR/tISR UIM because of its relatively tight binding to the Ub hydrophobic patch (K_d_ = 22 μM)^22,23^ and we predicted that a shorter linker (2 amino acids versus 5 with tIVR or tISR) could be used to connect the UQ1^UBA^ C-terminus with the N-terminus of IsoT^Buz^. The shorter linker was expected to reduce the entropic cost of complex formation. We conjugated Atto532 to UQ1^UBA^ G573C in tUI (Fig. 1a, lower right panel); titration with free Ub revealed a 3-fold increase in fluorescence and a remarkably low K_d_ of 66 pM (Fig. 1d,e). In an independent experiment, we measured the association and dissociation rate constants for the Atto532-tUI–Ub complex from which we calculated a K_d_ of 56.4 ± 1.2 pM (Supplementary Fig. S4), which is in excellent agreement with the results from titrations done at equilibrium. To our knowledge, this very high affinity is unprecedented for binding to a single Ub. Atto532-tUI showed exceptional selectivity against Ub C-terminal conjugates, having a binding preference for free Ub over Ub-GB1 of >10^6^. With respect to discrimination of Ub from UbL proteins, Atto532-tUI has 120-fold higher affinity for Ub than Nedd8. The substantially higher selection against Nedd8 seen with the tIVR and tISR sensors can be attributed to the Ruz domain, which is absent in tUI. Nonetheless, as described below, tUI’s selectivity coupled with its exceptional affinity make it the first choice for most *in vitro* applications.

The results above demonstrated the high selectivity of sensors engineered to bind Ub that has a free C-terminus. In cells, multiple forms of Ub can fulfill that criterion; most notably, the proximal Ub in a “free” or “unanchored” polyUb chain would have its C-terminal amino acids available for binding by the sensor Buz domain. Inspection of the sites on Ub used for Ub–Ub linkages in polyUb (i.e., Ub’s seven lysine ε-amines plus the α-amine on M1) showed that they are not occluded when bound by the various UBDs used in the sensors (Fig. 1a). These models are consistent with studies that have shown little or no discrimination by individual UBA, UIM, or Buz domains for binding Ub in chains with different Ub–Ub linkages^16,22,23^. Accordingly, competition by Ub_2_ or Ub_4_ linked through M1, K6, K29, K33, K48, or K63 revealed little difference among them or from free monoUb in binding to tIVR (Fig. 1e,f). Although not tested directly, based on previous findings of linkage-independent binding of UBA domains^22,23^ and the small tUI footprint modeled onto Ub (Fig. 1a), we anticipate similar linkage-independent binding by tUI to unanchored polyUb.

Another modification to Ub that potentially could affect detection by the sensors is phosphorylation. Multiple serine and threonine phosphosites have been found on Ub^24^; as with Ub lysines, most of these do not show overlap with the surfaces bound by our sensor UBDs (Fig. 1a). Ub phosphorylation on S65, which has functions in Ub-mediated signaling and mitophagy in particular, has been the most intensively studied phosphoUb species^25–28^. When we titrated Atto532-tUI with Ub(pS65) we determined a K_d_ of 262 pM (Fig. 1d,e), indicating 4-fold weaker binding than with Ub. Wauer et al. have shown by NMR that Ub(pS65) exists in an equilibrium between two principal conformers^29^. The major form (~70%) is essentially like unmodified Ub, whereas in the minor conformer (~30%) movement of the β5 strand retracts the normally-extended C-terminal residues into the body of Ub and displaces a key component of the hydrophobic patch. Because interactions with Buz and UBA domains are likely to be disrupted in the minor Ub(pS65) conformer, we expect that tUI would bind tightly only to the major conformer; thus, the Ub versus Ub(pS65) K_d_ difference could in part reflect the equilibrium where only a fraction of Ub(pS65) is in the binding-competent conformation.

### Real-time deubiquitination assays with label-free substrates

Quantitative activity assays are essential to understand the regulations and specificities of DUBs. Typically physiological substrates of DUBs are not used in quantitative assays due to limited availability and lack of good methods to quantify products. Moreover, real-time monitoring of activity — the preferred approach to high-precision kinetics studies — has been virtually impossible with physiological DUB substrates. For these reasons, artificial Ub derivatives such as Ub-(7-amido-4-methylcoumarin) or Ub–protein conjugates (e.g., diUb) with pairs of fluorophores that enable FRET-based assays are used^30,31^. Our free Ub sensors now make it possible to develop less restrictive DUB assays that could employ virtually any Ub conjugate as a substrate. As an example, we used Atto532-tIVR to monitor DUB-catalyzed release of free Ub (or unanchored polyUb) in a continuous fluorometric assay. For a model substrate, we used K48-linked Ub_5_ conjugated to the N-terminus of ovomucoid first domain (OM); to provide a second means to assay reactions, the OM moiety was modified with Lucifer Yellow dye (LY) to facilitate detection after SDS-PAGE^32^. For the DUB, we used human OTUB1, which selectively cleaves K48 Ub–Ub isopeptide linkages^33^ and is activated allosterically by interaction with certain E2 enzymes^34^. Without OTUB1, Ub_5_OM(LY) and Atto532-tIVR showed no fluorescence change, whereas enzyme addition initiated a fluorescence increase indicating release of free (poly)Ub (Fig. 2). Furthermore, in agreement with the report by Wiener et al.^34^, addition of UbcH5c stimulated the deubiquitination activity. SDS-PAGE of samples from the reaction mixtures confirmed the results from the sensor (Supplementary Fig. S5). Because OTUB1 is selective for K48-linked polyUb, Ub_1_-OM(LY) accumulated upon OTUB1 digestion, even in the presence of UbcH5c (lane 6, Supplementary Fig. S5a-c). This remaining conjugated Ub could be cleaved by the non-specific DUB Usp2cc (lane 7, Supplementary Fig. S5a-c). The release of free Ub determined in real-time using the sensor agreed with the amounts calculated by quantifying the LY-labeled gel bands (Supplementary Fig. S5d).

**Fig. 2.**
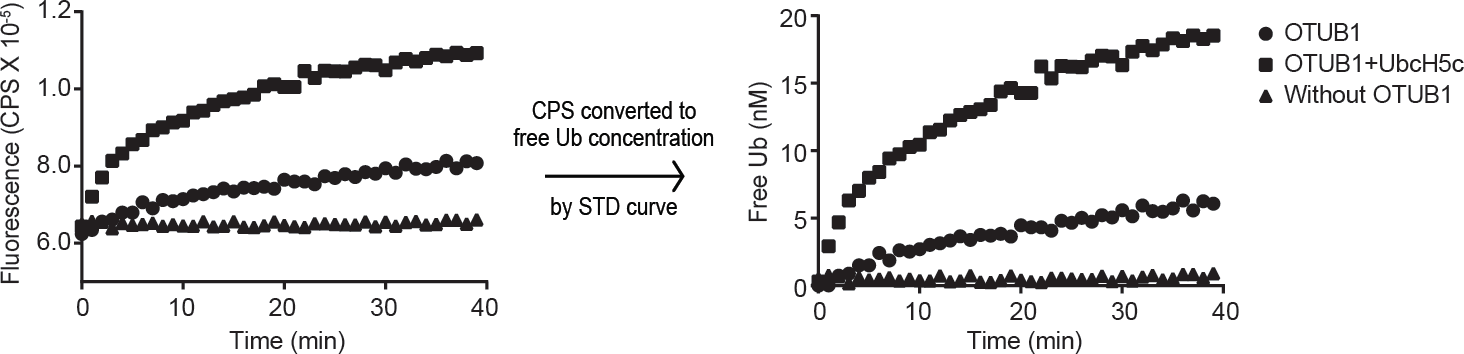
Quantitative, real-time DUB activity assays with a Ub sensor. K48-linked Ub_5_OM(LY) (10 nM), a mimic of a polyubiquitinated protein conjugate, was mixed with 5 μM OTUB1 with or without 20 μM UbcH5c at 25 °C in the presence of 2 nM Atto532-tIVR, and Atto532-tIVR fluorescence was monitored (left panel). A standard (STD) curve of Atto532-tIVR titrated with Ub (see Supplementary Fig. S5e) was used to convert the fluorescence intensity of Atto532-tIVR to free Ub concentration (right panel).

### Ub pool quantitation in cell lysates

We next developed methods that use the sensors to quantify cellular Ub pools. Our goal was to generate a workflow for routine measurements that would not depend on expensive, time-intensive steps requiring chromatography and mass spectrometry to separate and detect different Ub populations. The general approach, in which the sensor (e.g., Atto532-tUI) fluorescence is measured with and without addition of cell lysate or other sample, promised to be simple and direct. The main challenge was to develop conditions to prevent appearance of free Ub due to disassembly of conjugates by endogenous DUBs or from spontaneous hydrolysis of Ub thioesters.

Our strategy was to lyse cells and quickly inactivate endogenous DUBs and other proteases (see Methods) and then treat each sample in three ways to differentially convert Ub pools to the free-Ub form for measurement with the sensor (Fig. 3a). In order to measure endogenous free Ub without interference from chemically-labile Ub thioesters, samples were treated with hydrazine to rapidly and selectively convert all Ub thioesters into Ub C-terminal hydrazide; Ub-hydrazide is stable and, relative to free Ub, gives a negligible response with the sensor (Fig. 1b-e and Supplementary Fig. S6). Thus, sensor fluorescence of a hydrazine-treated sample will measure endogenous free Ub. With a second portion of the sample, β-mercaptoethanol was used to release Ub from Ub thioesters; measurement with the sensor then will report the sum of the endogenous free and thioester Ub pools^35^. A third portion was incubated with Usp2cc, a truncated DUB that can deubiquitinate virtually all forms of conjugated Ub^36^; when used in combination with a thiol reducing agent, all forms of Ub in the sample are converted to free Ub and the sensor readout will report the total Ub. By deducting the sum of the activated and free Ub from the total Ub, we can determine the amount of Ub that had been in (poly)Ub–proteins or other conjugates.

**Fig. 3.**
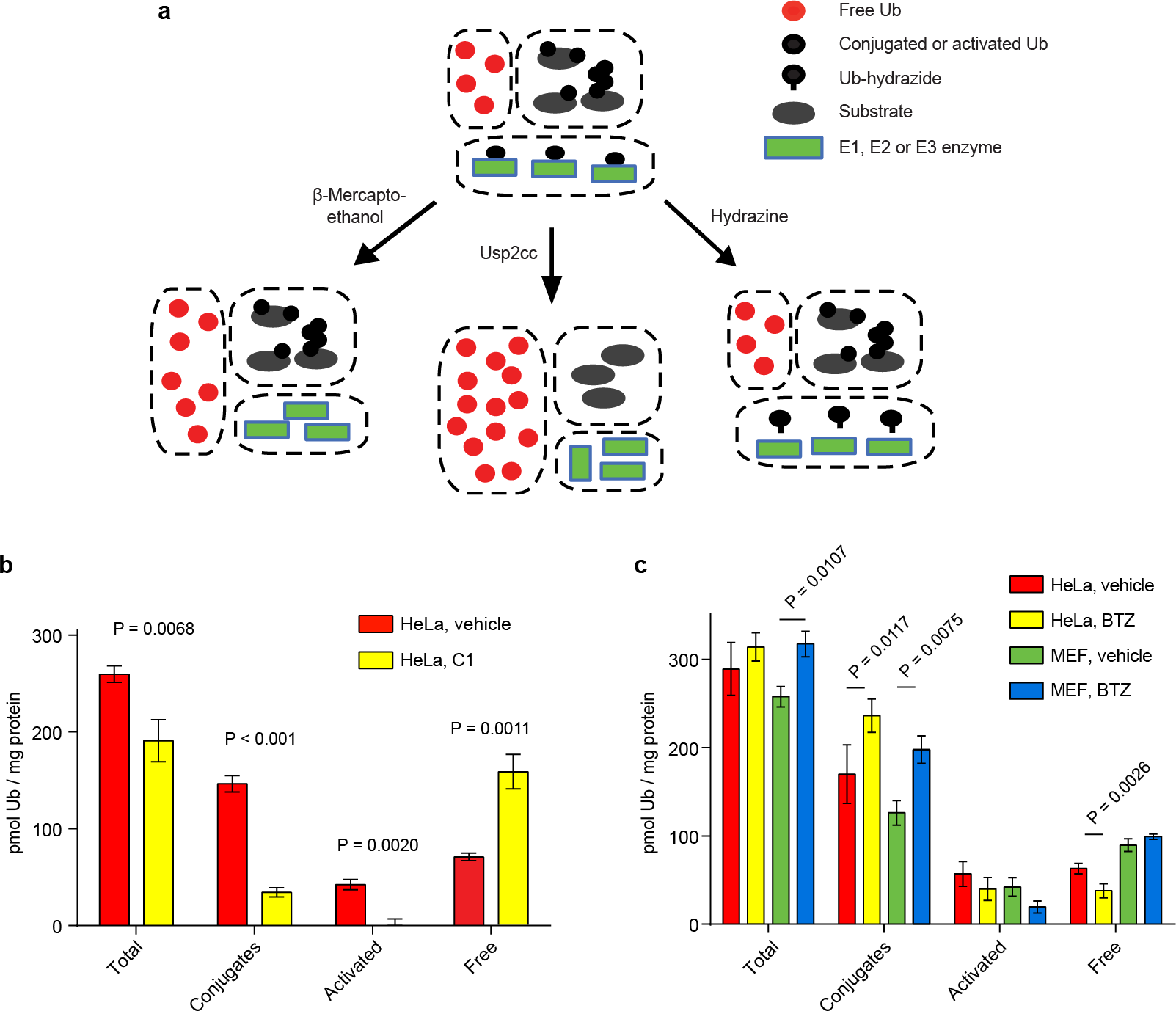
Effects of cellular stresses on Ub pools. **a**, Scheme used for the in-solution Ub pool measurements. **b,c**, Quantitation of Ub pools in lysates of indicated cell lines after treatment with vehicle (DMSO) or (**b**) E1 inhibitor, C1, at 10 μM or (**c**) proteasome inhibitor, BTZ, at 1 μM for 1 h. Statistical analyses by t-test (**b**) and ONE-WAY ANOVA with Bonferroni’s adjustment (**c**); error bars represent ± s.d. (n = 3).

These assays were first used to quantify the amounts of free, activated, and conjugated Ub in HeLa cells (Fig. 3); the results are in good agreement with other reports^15,37,38^. The sensor assays then were used with HeLa cells after treatment with inhibitors of the E1 Ub-activating enzyme or proteasome (Fig. 3b,c). As expected^39^, the E1 inhibitor Compound 1 (C1) dramatically increased free Ub with a concomitant loss of activated Ub and most Ub–protein conjugates (Fig. 3b). Conversely, proteasome inhibition by bortezomib (BTZ) promoted accumulation of Ub conjugates that reached a maximum at 1 h and then persisted through a 4 h treatment (Fig. 3c; Supplementary Fig. S7). The conjugate increase was accompanied by a modest depletion of activated Ub and a two-fold decrease in free Ub, presumably due to impaired proteasome-dependent recycling of Ub from conjugates. Different from this result, proteasome-inhibited MEF cells exhibited little change in free Ub levels, even though conjugated Ub increased 50%; instead, the total amount of Ub increased, likely due to increased expression of Ub genes^40^.

### Quantitation of endogenous free Ub in fixed cells

We realized that Atto532-tUI, with its high affinity and specificity for free Ub and conjugated fluorophore, could be an ideal tool to localize endogenous, intracellular free Ub. Initially, fixed cells were stained directly with Atto532-tUI, but high background fluorescence, most likely from nonspecific binding by the fluorophore, reduced sensitivity (data not shown). Therefore, we used instead a hemagglutinin-tagged version of tUI (HA-tUI) followed by detection with an anti-HA antibody. We confirmed the specificity of HA-tUI for free Ub by performing control experiments where fixed cells were incubated with HA-tUI together with excess free Ub. The fluorescence observed in these competition experiments was negligible, suggesting that the staining with HA-tUI is specific for cellular Ub (Supplementary Fig. S8). A diffuse intracellular distribution of free Ub was expected based on its small size (8.6 kDa) and negligible self-association^41^. Staining with HA-tUI was observed throughout the cytoplasm and nucleus in HeLa, U2OS, MEF and RPE1 cell lines (Fig. 4a). Compared to K48-linked chains and mono-and polyubiquitylated proteins, free Ub staining is evenly distributed through the whole cell (Supplementary Fig. S9). GFP-Ub that had been mutated to prevent its conjugation to other proteins was similarly diffuse when expressed in mammalian cells^13^.

**Fig. 4.**
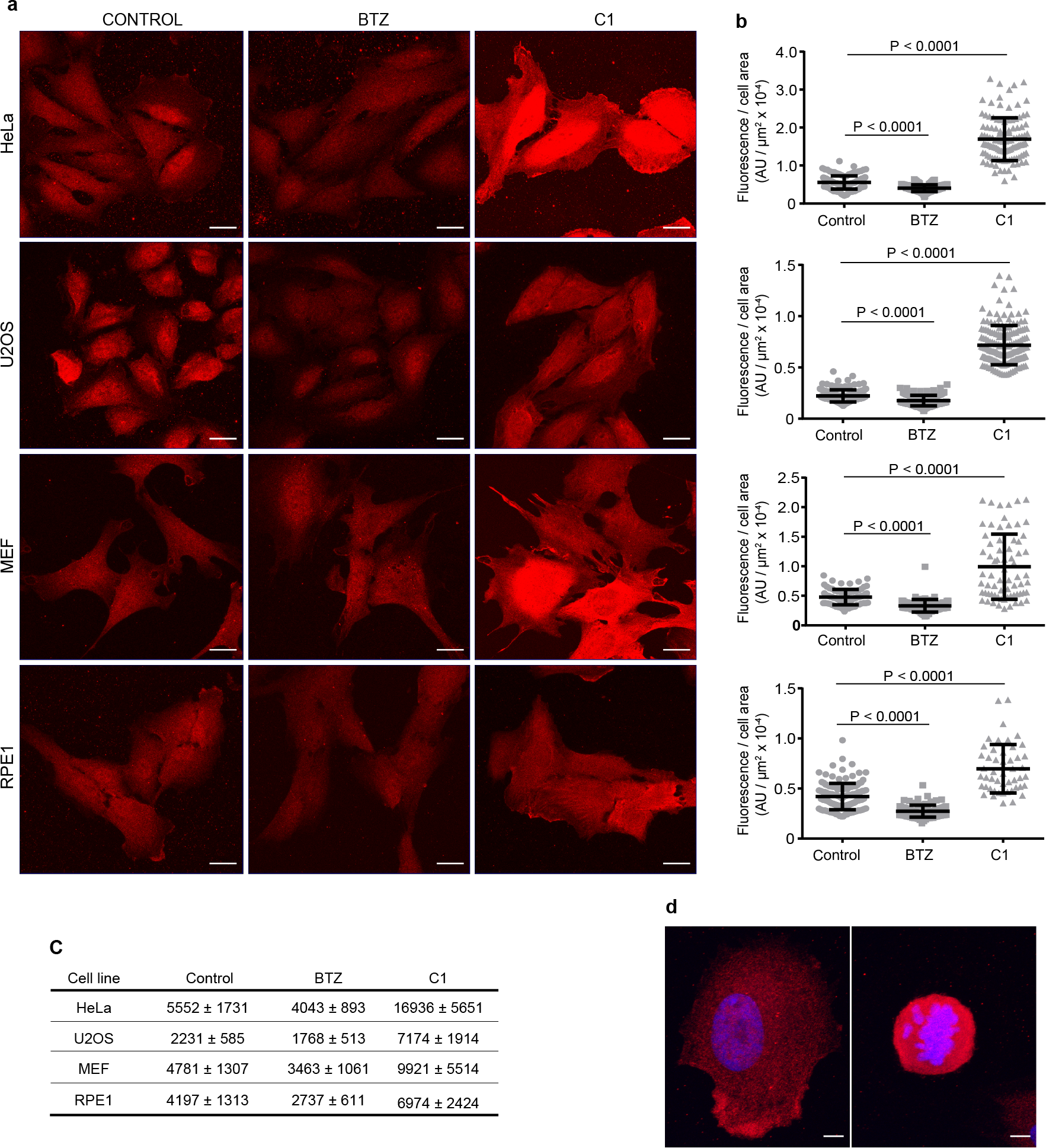
Free Ub staining in HeLa, U2OS, MEF and RPE1 cells. **a**, Maximum projection images of free Ub staining with HA-tUI in HeLa, U2OS, MEF and RPE1 cells after 1 h incubation with 1 μM proteasome inhibitor, BTZ or 10 μM E1 inhibitor, C1. Scale bars, 20 μm. **b**, Mean fluorescence for HeLa, U2OS, MEF, and RPE1 cells after 1 h incubation with 1 μM proteasome inhibitor or 10 μM E1 inhibitor. AU, arbitrary units. Cells analyzed per condition: HeLa, control n = 131, BTZ n = 125, C1 n = 116; U2OS, control n = 161, BTZ n = 170, C1 n = 175; MEF, control n = 80, BTZ n = 79, C1 n = 69; RPE1, control n = 133, BTZ n = 87, C1 n = 49. Bars show mean ± s.d. Statistical analyses by unpaired Student’s *t*-test with Welch's correction where appropriate. **c**,Relative free Ub from staining (mean fluorescence ± s.d.) of untreated cells or after proteasome or E1 inhibition. **d**,Representative interphase (left panel) and mitotic (right panel) RPE-1 cells stained with HA-tUI (red) and DAPI (blue). Intensity measurements from 3D reconstructions employed Imaris software. Total cell fluorescence (arbitrary units; mean ± s.d., n ≥ 3) for interphase and mitotic RPE1 cells were 23.7 ± 2.5 and 38.4 ± 3.5, respectively, whereas fluorescence per unit volume was 134.1 ± 8.6 for interphase RPE1 cells and 463.2 ± 70.6 for mitotic RPE1 cells. Scale bars, 5 μm.

Staining with HA-tUI offers an alternative to solution-based assays to monitor changes in free Ub during growth or in response to different stresses. After proteasome inhibition by incubation with BTZ for 1 h, HeLa, U2OS, MEF and RPE1 cells showed decreased staining with HA-tUI based on their 1.3 to 1.5-fold lower anti-HA antibody fluorescence relative to control (i.e., vehicle only) cells. As expected, E1 inhibition increased the free Ub (2-fold in MEF and RPE cells, and 3-fold in HeLa and U2OS cells) (Fig. 4b,c). Although proteasome inhibition decreased free Ub staining in all four cell lines, the intracellular distributions of free Ub appeared unaffected. For RPE1 cells, ratios of cytoplasmic to nuclear staining were not changed significantly by incubation with BTZ (control cells, 0.85 ± 0.07, n = 5; BTZ, 0.89 ± 0.08, n = 4). In contrast to the results with BTZ, increased staining was observed after E1 inhibition and cell-to-cell variability was greater overall (Fig. 4a-c). The average changes in staining of HeLa cells are consistent with the in-solution assays, whereas for MEF cells the staining showed ~25% less free Ub after proteasome inhibition than was determined from the in-solution assays (Figs. 3c and 4a-c). Possibly, because hydrazine treatment was not used with the imaged cells, some of the free Ub detected by staining could have originated from spontaneous hydrolysis of the Ub-thioester pool, thereby inflating the “free” Ub and confounding direct comparison of the two assays. The staining with tUI also revealed cell-cycle dependent differences in free Ub levels. RPE1 cells undergoing mitosis showed a 1.6-fold increase in tUI staining compared with interphase cells (Fig. 4d). The increase in free Ub may be due, at least in part, to the large-scale deubiquitination of H2A histones observed for mitotic cells^42^.

## Discussion

Individually, most UBDs have only modest affinity for Ub and are typically found together with other binding domains (and sometimes additional UBDs) to promote binding to specific types of polyUb or Ub-protein conjugates. From genetic fusions of multiple UBDs, we have engineered new proteins with specificity and extremely high affinity for monomeric free Ub. Our basic strategy was to use the Buz domain to direct binding to Ub with an unconjugated C-terminus, and to increase affinity with one or two additional UBDs that interact with Ub on non-overlapping surfaces. A critical aspect of the design strategy was to maximize the effect from avidity. This was achieved by having multiple UBDs bind simultaneously and by minimizing the entropy lost upon complex formation through careful selection of peptides linking the UBDs. Avid binding can boost affinity by combining the contributions of individual binding domains in a complex assembled from a multivalent ligand and a corresponding multivalent binder. The Gibbs free energy for binding overall (ΔG_total_) can be approximated as the sum of the individual binding domain (BD) interaction free energies (ΔG_BD1_, ΔG_BD2_, etc.) plus the unfavorable free energy due to reduction in entropy (predominantly, losses in translational and rotational entropy) from having all the binding domains linked together (ΔG_S_)^43^: 

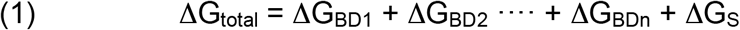

To achieve “perfect” avid binding and maximize affinity, ΔG_S_ should be close to zero. For Ub binding by a tandem-UBD (tUBD) protein such as tIVR, we can write: 

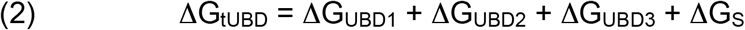

ΔG_UBD_ can be calculated for each UBD based on the K_d_ values reported for Ub binding by the individual IsoT^Buz^, Vps27^UIM^, and Rabex5^Ruz^ domains (i.e., −7.53, −5.25, and −6.62 kcal·mol^−1^, respectively); similarly, ΔG_tIVR_ = −10.52 kcal·mol^−1^ can be calculated from the tIVR–Ub K_d_ of 19.3 nM (Fig. 1e). Thus, although avidity promotes tight binding by tIVR, it’s at the cost of ΔG_S_, which is (−10.52) – (−19.40) = 8.88 kcal·mol^−1^. This substantial penalty reduced tIVR affinity by >10^6^ from a theoretical K_d_ of 5.9 × 10^−15^ M.

Remarkably, tUI’s affinity for Ub (K_i_ = 194 pM; Supplementary Fig. S10) is only 3-times weaker than what would be predicted for *perfect* avid binding by the combination of the UQ1^UBA^ domain (K_d_ = 22 μM)^22,23^ and the Buz domain (K_d_ = 3 μM)^16^. Contributing to the much greater efficiency of avidity seen with tUI relative to tIVR is that tUI has only one linker peptide (versus two in tIVR), and its linker is very short (Supplementary Fig. S1). Thus, a likely route to increase affinity for tIVR or tISR is further optimization of the interdomain linkers. On the other hand, a large increase in affinity by tUI would have to come from tighter-binding versions of one or both of its UBDs, as the linker peptide already is nearly optimum.

There are several ways in which our tUBD binders for free Ub could be improved in affinity or for specific applications. Affinity might be increased, as noted above, with either modified linkers or alternative UBDs. Our designs have utilized only a few of more than 20 types of UBDs^44,45^; moreover, other Ub binding proteins such as catalytically-inactive DUBs might be used as components of tUBD-type fusion constructs^46^. Particularly intriguing is the prospect of tailoring recognition by incorporation of a UBD that would select either for or against specific modifications on the Ub. For example, although tIVR binds equally well to free polyUb chains of different Ub–Ub linkage types (Fig. 1f), alternatives to the Ruz or UIM domain could be developed in which steric clash prevents binding to Ub conjugated at particular lysine(s). Similarly, UBD (or linker) modifications might be made to introduce selectivity for phosphoUb derivatives. In addition, alternative fluorophores and attachment sites can be explored further to improve sensor sensitivity and dynamic range.

As multiple DUBs have been implicated in human disease^47^, the sensors have potential to facilitate drug development. Although some DUB inhibitors have been tested clinically^47^, inhibitor development remains limited by difficulties in establishing high-throughput screens that can employ physiological DUB substrates. Important innovations of the sensor-based real-time assays are that virtually any Ub conjugate can be used as the substrate, and it is unnecessary to have the substrate labeled.

Other likely applications of the free Ub sensors are quantitations of Ub pools using the protocols we have described for in-solution assays or cell staining. Many neurological disorders disrupt Ub homeostasis and show aggregates of ubiquitinated proteins or a general depletion of free Ub. Staining by tUI offers a unique approach to examine, for example, neurons at the single-cell level to understand better the effects of genetic or environmental perturbations on Ub homeostasis and intracellular distribution. Additionally, extracellular free Ub has been suggested as a biomarker for trauma and disease^48^; here, the free Ub sensors can replace the less specific antibody-based assays typically used to quantify extracellular Ub. Ultimately, we envision that the free Ub binders and the fluorescent sensors developed from them will provide effective tools to capture, deplete, quantify, or visualize free Ub *in vitro* and in cells.

## Methods

### Materials and protein preparation

tIVR, tISR, tUI, Ub-GB1, and UbcH5c were cloned into pET28a and transformed into BL21-CodonPlus (DE3) E. coli cells for protein expression. Expression was induced by the addition of 0.4 mM IPTG to cells grown at 37 °C to OD_660nm_ = 0.6-0.8, and then growth was continued at 25 °C for 8 h. The cells were harvested by centrifugation at 3,200 × *g*, resuspended in ice-cold Buffer A (20 mM sodium phosphate, pH 7.4, 500 mM NaCl, 10 mM imidazole, and 10 mM β-mercaptoethanol), and lysed by sonication; the lysates were clarified by centrifugation for 30 min at 4 °C at 20,199 × *g*. A Histrap HP column (GE Healthcare, 17-5248-02) was used to purify the proteins from the lysates. Samples were applied to the column equilibrated with Buffer A, and after washing with 20 column volumes, bound proteins were eluted with a linear gradient to 500 mM imidazole in Buffer A. The proteins were further purified by gel filtration through a Superdex 75 column (GE Healthcare, 29-1487-21) eluted with pH 7.4 PBS and 1 mM DTT or 1 mM TCEP. Purity was confirmed by SDS-PAGE. Ub^49^, Nedd8^50^, Usp2cc^36^, and Ub_5_-OM(LY)^32^ were prepared as described. OTUB1 was a gift from C. Wolberger (Johns Hopkins University). K6, K11, K27, K29, K33, K48, and K63-linked Ub_2_ chains were purchased from UbiQ Bio (Amsterdam), Ub(pS65) was from BostonBiochem (Cambridge, MA), and M1-linked Ub_4_ was prepared as described^51^.

### Synthesis of Ub-hydrazide

Ub (1.5 mM) was incubated with 10 mM ATP, 10 mM MgCl_2_, 100 mM sodium 2-mercaptoethanesulfonate (MESNA; Fluka), and 100 nM mouse E1 in 20 mM HEPES (pH 8.0) for 3 h at 37 °C to form Ub-MESNA thioester (confirmed by mass spectrometry; see Supplementary Fig. S6b). The Ub-MESNA then was incubated in 300 mM aqueous hydrazine for 30 min at 37 °C to form Ub-hydrazide. The reaction product was diluted 25-fold with 50 mM ammonium acetate, adjusted to pH 4.5 (Buffer B), and purified by cation-exchange chromatography on a Mono S column (GE Healthcare, 17-0547-01). The column was washed with 20 volumes of Buffer B and eluted with a linear gradient of 0 – 1 M NaCl in the same buffer. The purified Ub-hydrazide was confirmed by mass spectrometry (Supplementary Fig. S6b).

### Fluorophore labeling

Sensor proteins were labeled at cysteine with fluorophore-maleimide dyes from ATTO-TEC GmbH (Atto dyes; see Supplementary Fig. S3), Molecular Probes (Alexa Fluor 488), or Anaspec (fluorescein). Fluorophore-maleimide dyes (1.5 to 5-fold molar excess) were incubated with 50 μM sensor in 50 mM HEPES, pH 7.4, 100 mM NaCl for 2 h at 25 °C. Excess dyes were quenched by incubation with 10 mM β-mercaptoethanol for 10 min at 25 °C. To remove excess dyes, the reaction product was bound to Ni-NTA resin (Thermo Fisher) equilibrated with Buffer A, the resin was washed 5 times with the Histrap binding buffer, and sensor proteins were eluted with Histrap elution buffer. Labeling was confirmed by SDS-PAGE and then scanning the gel for fluorescence using a Typhoon FLA 9500 (GE Healthcare Life Sciences). Degree of labeling (DOL) and concentrations of the labeled proteins were calculated by the equations below. 

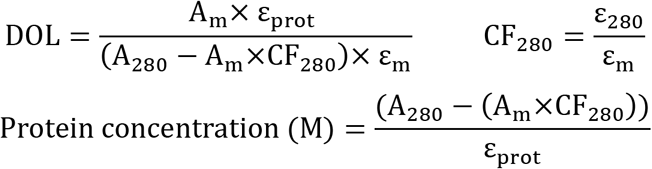

In the equations, A_m_ represents the absorbance at the dye absorption maximum, A_280_ is absorbance at 280 nm of the labeled protein, ε_prot_ is the extinction coefficients at 280 nm of the protein, ε_280_ is the extinction coefficient at 280 nm of the dye alone, ε_m_ is the extinction coefficient at the absorption maximum of the dye, and CF_280_ is the correction factor at 280 nm.

### Binding assays

All binding assays were done in PBS buffer, pH 7.4, supplemented with 0.05% Brij35 and either 0.2 mg/ml ovalbumin or GB1 protein, and either 1 mM DTT or 1 mM TCEP. A FluoroMax-4 spectrofluorimeter (HORIBA Scientific) was used to measure fluorescence intensity in the binding assays. K_d_ and K_i_ values were calculated by fitting with a single-site binding model^52^ using Prism 6 (GraphPad Software). Because Atto532-tUI has exceptionally high affinity for Ub, 10 pM Atto532-tUI was used in the binding assays to keep its concentration below the K_d_ for Ub. To improve detection of the fluorescence from this low concentration of Atto532-tUI, we increased the assay volume to 2.7 ml. The stock Ub titrated into the solution was ≤ 1.5% of the total volume.

### Stopped flow kinetics

To determine the k_off_ and k_on_ rates of Atto532-tUI with free Ub, rapid kinetics were monitored by fluorescence using a MOS-500 spectrometer equipped with a SFM-4000 mixer (Bio-Logic Science Instruments) maintained at 25 °C. The excitation wavelength was set to 530 nm with a 5 nm bandwidth, and the fluorescence emission was detected with a 540-620 nm bandpass filter.

### Real-time DUB assays

A Fluoromax-4 spectrofluorimeter (HORIBA Scientific) was used to monitor fluorescence of samples (45 μL) in ultramicro quartz cuvettes (Hellma) at 25 °C. The buffer was PBS, pH 7.4, with 0.05% Brij35, 0.2 mg/ml ovalbumin, and 1 mM DTT. A standard curve was generated to convert change in sensor fluorescence to the corresponding free Ub concentrations; e.g., 2 nM Atto532-tIVR was titrated with 2, 3, 6, 10, 18, and 32 nM Ub, and the binding curve was fit as described above. Fluorescent intensities from real-time DUB assays then were converted to free Ub concentrations using the fitted binding equation. For deubiquitination of Ub_5_-OM(LY), results were confirmed by SDS-PAGE of samples from the reaction mixtures and fluorescence imaging of the gel with a Typhoon FLA 9500 laser scanner (GE Healthcare Life Sciences).

### In-solution Ub pool assays

#### Sample preparation

Cells were lysed in 100 mM MOPS, pH 6.0, 8 M urea, 20 mM NEM, and EDTA-free Complete protease inhibitors (Roche) by sonication and then centrifuged at 15,800 × *g*. Total protein in the clarified extract was measured using the bicinchoninic acid (BCA) assay before being divided into three fractions for treatment with Usp2cc, β-mercaptoethanol, or hydrazine. The fraction to be treated with Usp2cc was diluted with digestion buffer (25 mM HEPES, pH 7.5, 140 mM NaCl, and 10 mM DTT) to reduce the urea to less than 2 M; to this, Usp2cc was added at a 1:10 (Usp2cc:total protein) ratio and incubated at 37 °C for 1 h. To another fraction of the extract, 100 mM CHES, pH 9, containing 150 mM β-mercaptoethanol was added and incubated at 37 °C for 1 h. The third fraction was incubated at 37 °C for 1 h with freshly-made 200 mM hydrazine-HCl, pH 8.5. These samples were then diluted using PBS and 0.2 mg/ml ovalbumin to insure that [Ub] was within the linear range of the assay (e.g., with Att532-tUI, from 2-60 nM). Dilution was also performed to reduce the concentrations of urea (<0.2 M), β-mercaptoethanol (<20 mM), hydrazine (<20 mM), which otherwise can interfere with binding by the sensor.

#### In-solution high-throughput assay to measure free Ub

Microplates (384-well SensoPlate Plus, Greiner Bio-One 781856) were used to measure free Ub concentrations of unknown samples in a high-throughput format. The plates were passivated by sequential treatment with 1% Hellmanex detergent, 1 M KOH, and then 2% 1,7-dichloro-octamethyltetrasiloxane (Sigma-Aldrich) diluted in heptane, where each step was a 30 min soak followed by extensive washes with distilled water and finally air-drying. Typically, 24 μl of a master mix containing 50 nM Atto532-tUI and assay buffer (200 mM sodium phosphate, pH 7.5, 50 mM NaCl, 2 mM DTT, 0.05% Brij35, and 0.2 mg/ml ovalbumin) was added to wells of the passivated 384-well plate. Then, to one set of wells, Ub standards were added and the remaining wells were used for samples (6 μl). Fluorescence intensities were quantified with a Typhoon FLA 9500 laser scanner (GE Healthcare). The Ub concentrations of unknown samples were determined by interpolating the fluorescent signals on standard curves generated by titration with a standard of free Ub.

### Sample preparation for microscopy

HeLa, U2OS, and MEF cells were cultured in DMEM (Gibco) supplemented with 10% (v/v) FBS and 100 U/mL penicillin and 100 ug/mL streptomycin (Hyclone). RPE1 cells were cultured in DMEM/Ham’s F-12, 50/50 mix (Corning). Cells were incubated for 1 hour with 1 μM bortezomib (Ubiquitin-Proteasome Biotechnologies), 10 μM E1 inhibitor (Compound 1; provided by Takeda Oncology, Cambridge, MA) or vehicle alone (0.1% DMSO). Each experiment was performed a minimum of two times.

We fixed cells at <80% confluence with 4% paraformaldehyde in PBS for 30 min at 37 °C, permeabilized them with 0.1% Triton X-100 for 10 min at room temperature and blocked for 1 hour with 5% BSA and 0.5% Tween 20 in PBS. Cells were stained for free Ub with 100 nM tUI-HA diluted in blocking solution for 30 min at room temperature. As a negative control, the sensor was pre-incubated for 5 min at room temperature with 100 μM Ub in blocking solution before addition to the samples. Next, cells were incubated overnight at 4 °C with anti-HA antibody (Sigma-Aldrich clone HA-7 or Bethyl Laboratories A190-108A; 1:1000 dilution), stained with Alexa Fluor 568-conjugated goat anti-mouse IgG (Thermo Fisher; 1:500 dilution), and mounted on slides using ProLong Diamond Antifade medium (Thermo Fisher). Some coverslips were also stained with anti-Ub (clone FK2, Biomol; 1:1,000 dilution) or anti-K48Ub (clone Apu2, Millipore; 1:200 dilution) primary antibodies and, subsequently, with Alexa Fluor 568-conjugated goat anti-mouse and Alexa Fluor 488-conjugated goat anti-rabbit (Thermo Fisher; 1:400 dilution) secondary antibodies. The HCS CellMask dye (Thermo Fisher) was added to the cells for half hour as a marker of cell boundaries for high-content fluorescence intensity-based measurements.

### Microscopy and image analysis

Cells were imaged using a Zeiss LSM 880 confocal microscope with C-Apochromat 40X/1.20 W or Plan-Apochromat 63x/1.40 Oil DIC M27 objectives. Z-stack images were acquired with ZEN Black software (Version 14.0.9.201) at 0.746 μm intervals. ImageJ 1.51h (NIH) was used to perform maximum intensity projections of z-sections and to calculate cell mean fluorescence intensity values; cell contours were drawn using HCS CellMask dye as a reference, and nuclei were identified with DAPI stain. Autofluorescence intensities recorded from unstained cells were subtracted from the tUI-fluorescence.

Three-dimensional reconstructions of RPE-1 cells were obtained with Imaris software (version 9.1.1, Bitplane AG) from serial z-sections acquired at 0.242 μm increments. The Imaris surface creation tool was used to generate volume renderings and to quantify tUI fluorescence intensities in both interphase and mitotic cells.

### Statistical Analysis

Statistical calculations were performed with GraphPad Prism software and are described in the relevant figure legends. *P* values less than 0.05 were considered significant.

## Supporting information

Supplement for Choi et al.

## Acknowledgements

We thank O. Peersen for assistance with the rapid-kinetics experiments and use of the Bio-Logic stopped-flow spectrofluorometer, C. Wolberger for OTUB1 protein, B. Brasher for phosphoubiquitin, and R. Handa for use of the Imaris image analysis software. This research was supported by NIH-NIGMS grant R01 GM37666 (to R.E.C.) and NIH-NIEHS grant R21 ES029150 (to R.E.C. and T.Y.).

## Author contributions

Y.C. and R.E.C. conceived and designed the ubiquitin sensor reagents, and Y.C. produced the sensors and characterized them in vitro. S.B., T.Y., and R.E.C. conceived of the cell-based studies, which were done by S.B., L.P., and F.S. All authors contributed to writing or commenting on the manuscript.

