## Supplement for Choi et al. for "High-affinity free ubiquitin sensors as quantitative probes of ubiquitin homeostasis and deubiquitination"

Supplementary Figure S1

tIVR

MGSSHHHHHHSSGLVPRGSHMKQEVQAWDGEVRQVSKHAFSLKQLDNPARIPPSG
WKCSKCDMRENLWLNLTDGSILCGRRYFDGSGGNNHAVEHYRETGYPLAVKLGTITP
DGADVYSYDEDDMVLDPSLAEHLSHFGIDMLKMQKGSCAAAEE**A**ELDLKAAIQESLRE
AGGGSDLLCKKGCGYYGNPAWQGFCSKCWREEYHKARQK

tISR

MGSSHHHHHHSSGLVPRGSHMKQEVQAWDGEVRQVSKHAFSLKQLDNPARIPPSG
WKCSKCDMRENLWLNLTDGSILCGRRY**W**DGSGGNNHAVEHYRETGYPLAVKLGTIT
PDGADV**W**SYDEDDMVLDPSLAEHLSHFGIDMLKMQKGSCAAEE**A**EEQIAYAMQMSL
REAGGGSDLLCKKGCGYYGNPAWQGFCSKCWREE**A**HKA**A**QK

tUI

MPSSHHHHHHSSGLVPRGSHMGSTVRFQQQLEQLSAMGFLNREANLQALIATCGDIN
AAIERLLGSGSEVRQVSKHAFSLKQLDNPARIPPSGWKCSKCDMRENLWLNLTDGSIL
CGRRYFDGSGGNNHAVEHYRETGYPLAVKLGTITPDGADVYSYDEDDMVLDPSLAEH
LSHFGIDMLKMQK

tUI-HA

MPSSHHHHHHSSGLVPRGSHMGSTVRFQQQLEQLSAMGFLNREANLQALIATCGDIN
AAIERLLGSGSEVRQVSKHAFSLKQLDNPARIPPSGWKCSKCDMRENLWLNLTDGSIL
CGRRYFDGSGGNNHAVEHYRETGYPLAVKLGTITPDGADVYSYDEDDMVLDPSLAEH
LSHFGIDMLKMQKCGSGSGYPYDVPDYAS

**Fig. S1. Primary sequences of the free Ub sensors.** The bold, underlined residues indicate mutated residues, and the cysteines in red indicate fluorophore conjugation sites. The residues highlighted in cyan are linkers introduced to connect the domains, and the yellow and green highlighted sequences show peptides added to provide 6-His and HA-epitope tags, respectively.

Supplementary Figure S2

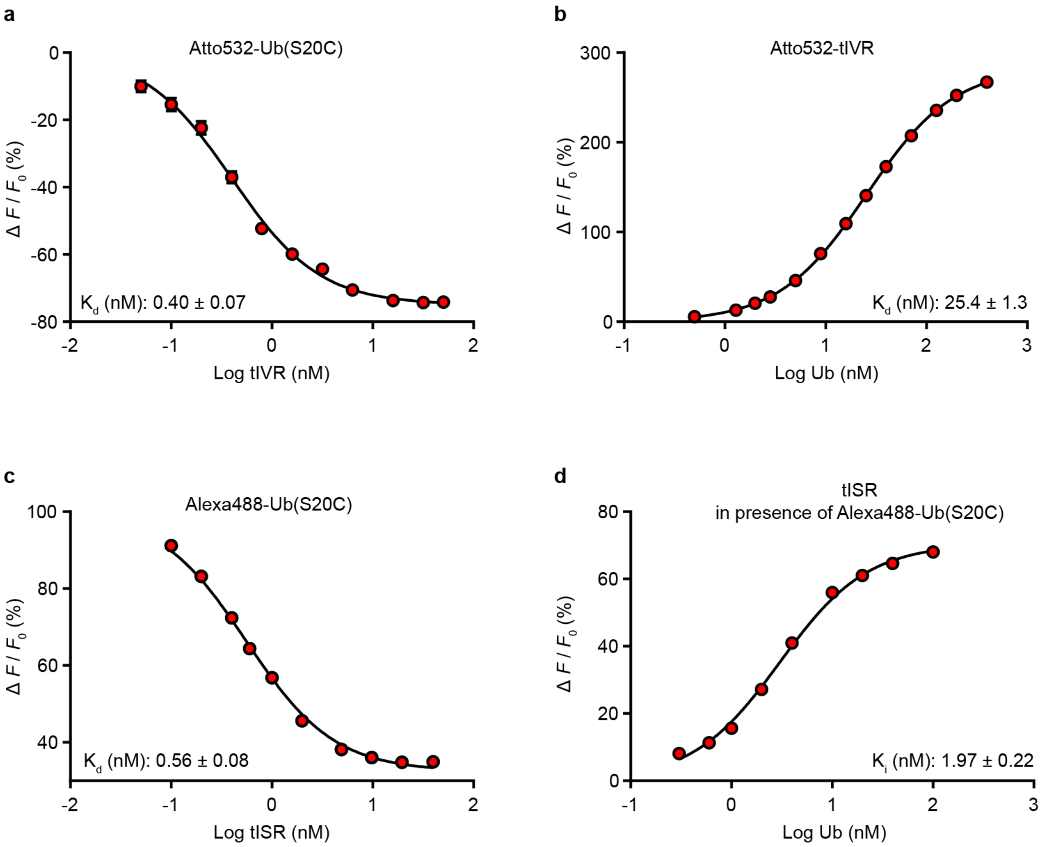


**Fig. S2. Sensor binding assays. a,** Atto532-Ub(S20C) titrated with tIVR. **b,** Atto532-tIVR titrated with Ub. **c,** Alexa488-Ub(S20C) titrated to tISR. **d,** tISR competition binding assay performed by titration of Ub in presence of Alexa488-Ub(S20C). In each titration, the fluorescence intensity change from ligand binding was measured. Error bars in **a**, which are mostly hidden by the point symbols, show standard deviations of the assay repeated two times; titrations in **b-d** were performed once.

Supplementary Figure S3

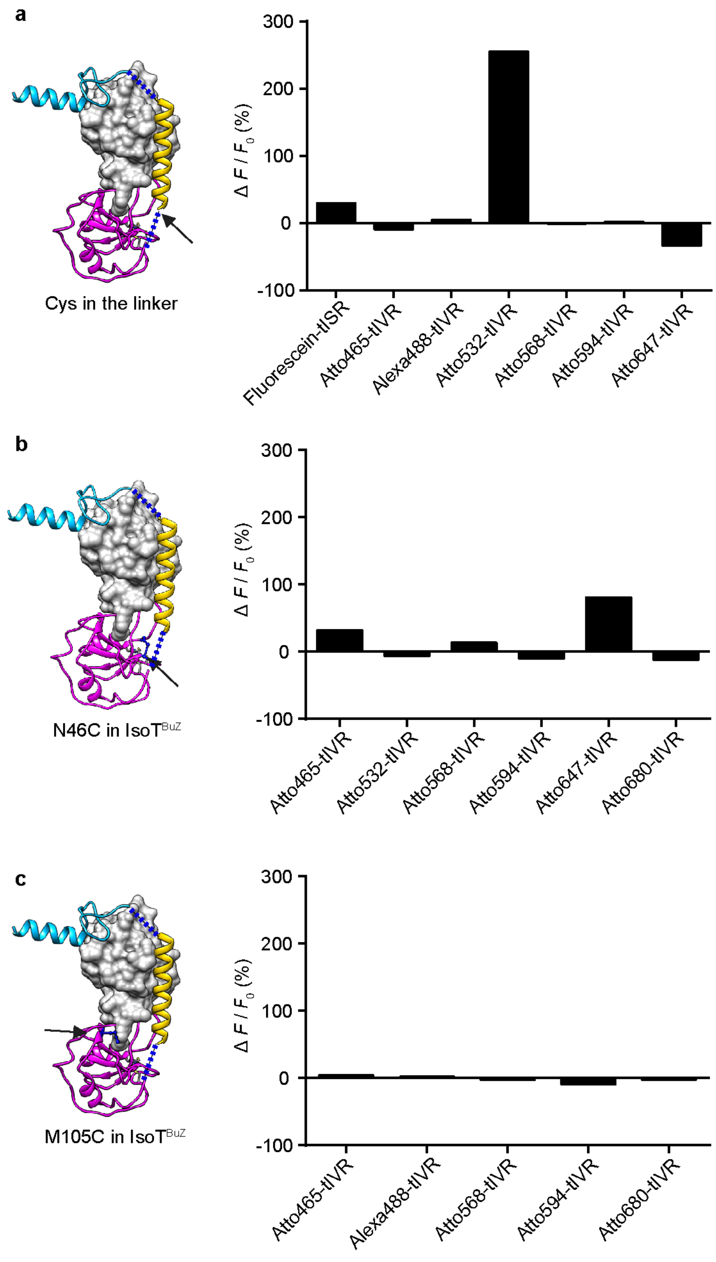


**Fig. S3. Screening for an optimal fluorescent dye and conjugation site on the tIVR sensor. a,** Cys residue in the linker connecting $IsoT^{Buz}$ and $Vps27^{UIM}$; **b**, N46C; and **c**, M105C in $IsoT^{Buz}$ were selected to conjugate various fluorescent dyes. In each panel, the arrow shows the conjugation site on the tIVR model and the histograms show performance with different fluorophores installed at that site. To determine which site and fluorescent dye gave the largest fluorescence change upon Ub binding, fluorescence intensities of the labeled sensors (2 nM) were measured with and without saturating Ub. $F_0$ is fluorescence intensity of the labeled sensors without Ub, and $\Delta F$ is the intensity change upon addition of 200 μM Ub.

Supplementary Figure S4

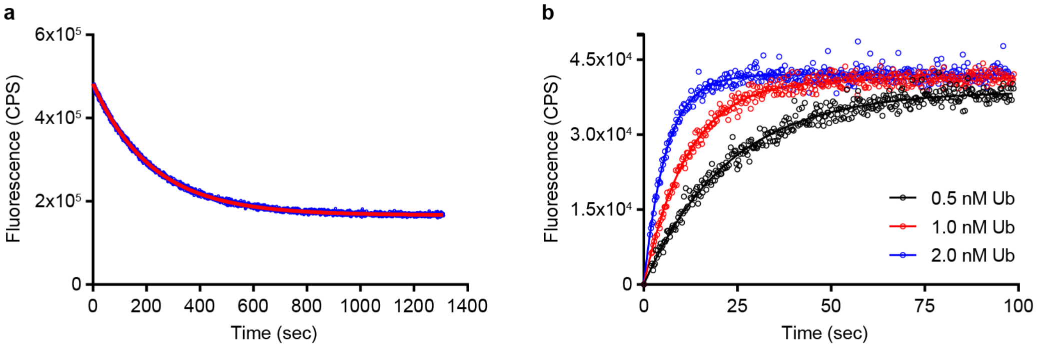


c

| Association ($M^{-1}$ $s^{-1}$) | Dissociation ($s^{-1}$) | $k_{off}$ / $k_{on}$ (pM) |
|---|---|---|
| 8.04 x $10^7$ ± 8.6 x $10^5$ | 0.00454 ± 0.00027 | 56.4 ± 1.2 |

**Fig. S4. Kinetics of Atto532-tUI binding to Ub. a,** To determine the dissociation rate of the Atto532-tUI•Ub complex, Atto532-tUI (1.0 nM) was pre-incubated at 25 °C with 1.0 nM Ub, and then excess unlabeled tUI (1 μM) was rapidly added and the fluorescence was measured every 0.25 s using a stopped-flow fluorimeter. The reaction was repeated three times, and the averaged data were fit with a single-exponential decay model to determine $k_{off}$. The experimental data (blue) are shown superimposed with the fitted curve (red). **b,** The change in fluorescence intensity of 50 pM Atto532-tUI immediately after addition of 0.5, 1.0, or 2.0 nM Ub was monitored at 25 °C. The fluorescence was measured every 0.25 s. From each curve, an observed rate ($k_{obs}$) was determined from the best-fitting exponential curve, and from these $k_{on}$ was determined from the equation $k_{obs} = k_{on} * [Ub] + k_{off}$. **c,** The experimentally determined association and dissociation rates for the Atto532-tUI•Ub complex are shown together with the $K_d$ calculated from their ratio.

Supplementary Figure S5

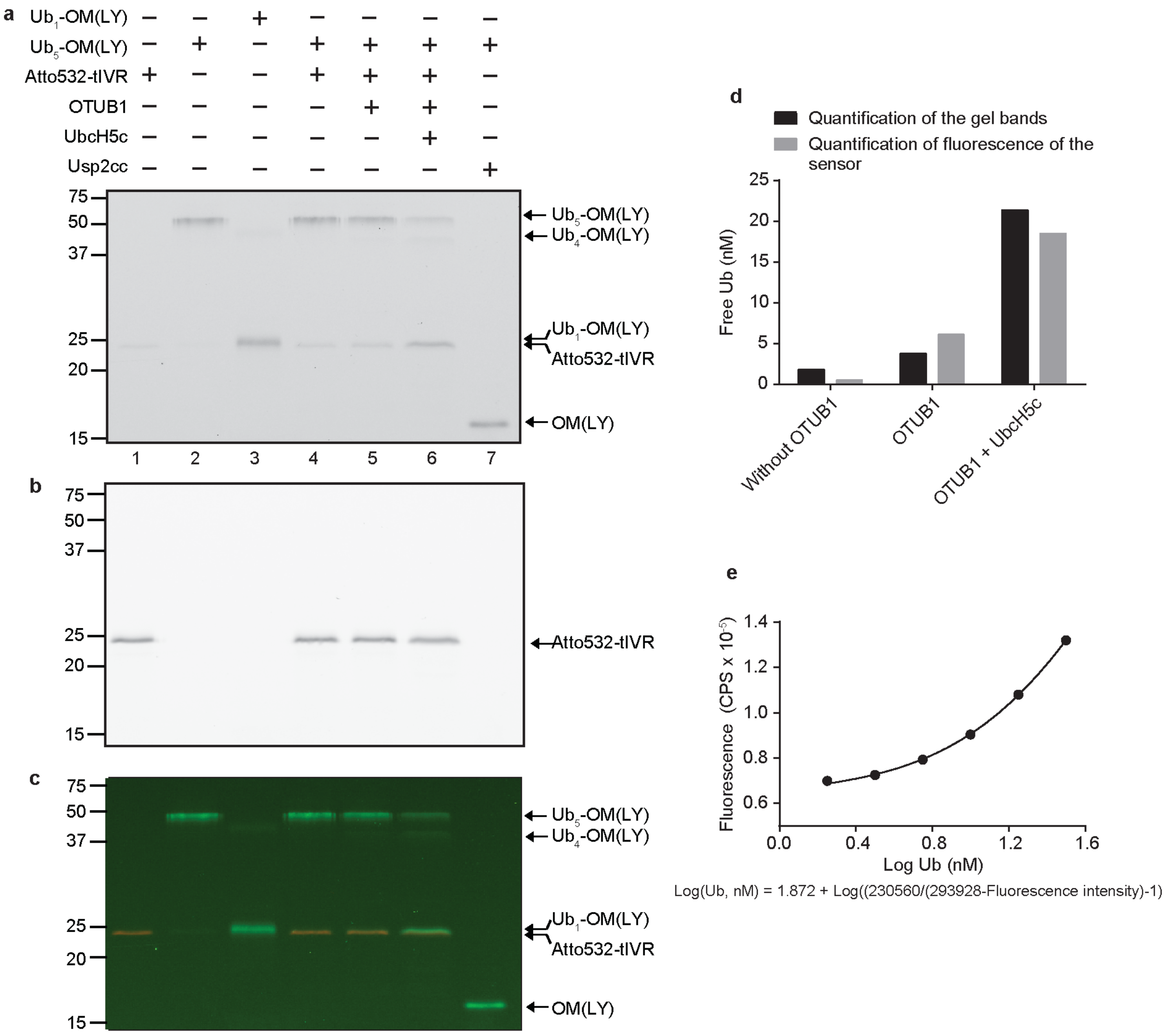


**Fig. S5. Analysis of the DUB reaction products by SDS-PAGE.** Samples from the end-points of the real-time DUB reactions in Fig. 2 were separated by SDS-PAGE together with the following control samples: Atto532-tIVR in lane 1, $Ub_5$-OM(LY) in lane 2, $Ub_1$-OM(LY) in lane 3, and the Usp2cc-digested product in lane 7. Fluorescent gel bands were detected with a Typhoon laser scanner using excitation at either **(a)** 488 nm, **(b)** 532 nm, or **(c)** both to visualize Lucifer Yellow-labeled ovomucoid protein (OM(LY)) or Atto532-labeled tIVR. **d,** The bar graph compares the Ub released in the DUB reactions in Fig. 2 quantified by either the protein band fluorescence (panel **a**) or the sensor fluorescence (Fig. 2). **e,** Atto532-tIVR (2 nM) was titrated with Ub (2 - 32 nM) and the data were fit with a 1:1 binding model as described in Methods. The fitted equation was used to convert fluorescence intensities from the real-time DUB reaction curves (Fig. 2, left panel) to free Ub concentrations (Fig. 2, right panel).

Supplementary Figure S6

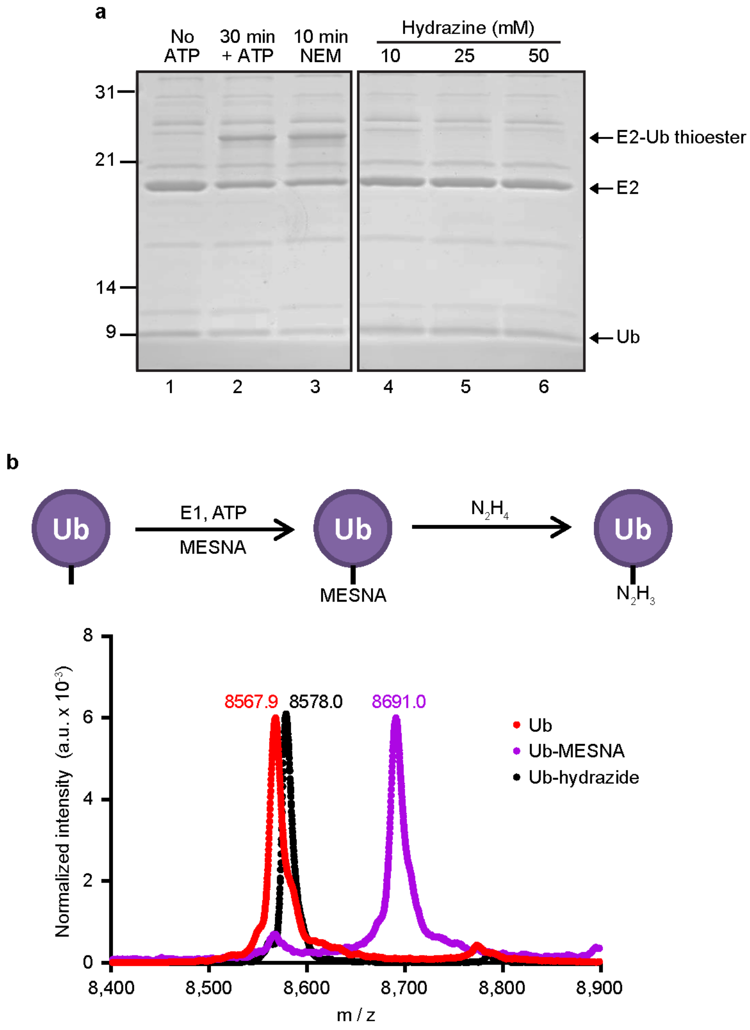


**Fig. S6. Generation of Ub-hydrazide from Ub thioesters. a,** Hydrazinolysis of thioester-linked Ub. Ub was incubated with E1 and E2 (UbcH5c) enzymes without (lane 1) or with (lane 2) ATP to form Ub~UbcH5c thioester. After treatment with NEM (lane 3) to inactivate enzymatic activities, hydrazine was added at the concentrations indicated; hydrazinolysis was complete under all three conditions. **b,** Upper panel shows the reaction scheme to generate Ub-hydrazide (see Methods) for Fig 1b-d titrations. Lower panel shows overlayed MALDI-TOF mass spectra of Ub and products (i.e., Ub–MESNA and Ub–hydrazide) from each step in the synthesis.

Supplementary Figure S7

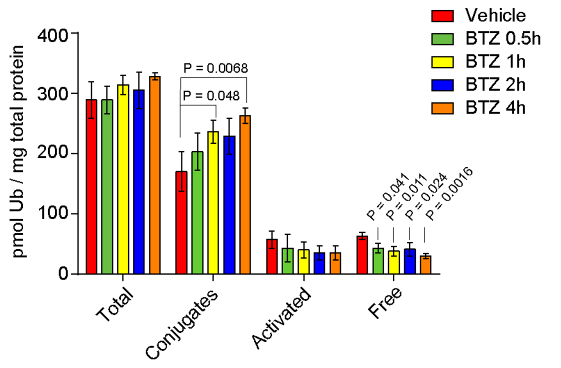


**Fig. S7. Ubiquitin pools in proteasome-inhibited HeLa cells do not change significantly after 1 h.** In-solution quantification of Ub pools in lysates after treatment with vehicle (DMSO) or the proteasome inhibitor, BTZ (1 µM). Statistical analysis was by one-way ANOVA with Bonferroni's adjustment; error bars represent ± s.d. (n = 3).

Supplementary Figure S8

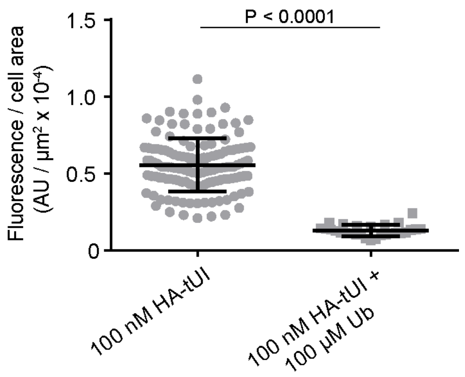


**Fig. S8. Competition by free Ub for cell staining by HA-tUI.** Quantitation of mean fluorescence in HeLa cells (maximum projection images) after incubation with HA-tUI with or without excess free Ub. For 100 nM HA-tUI, n = 131; for 100 nM HA-tUI + 100 μM Ub, n = 28. AU, arbitrary units; error bars indicate mean ± s.d. Statistical analysis used unpaired Student's *t*-test with Welch's correction.

Supplementary Figure S9

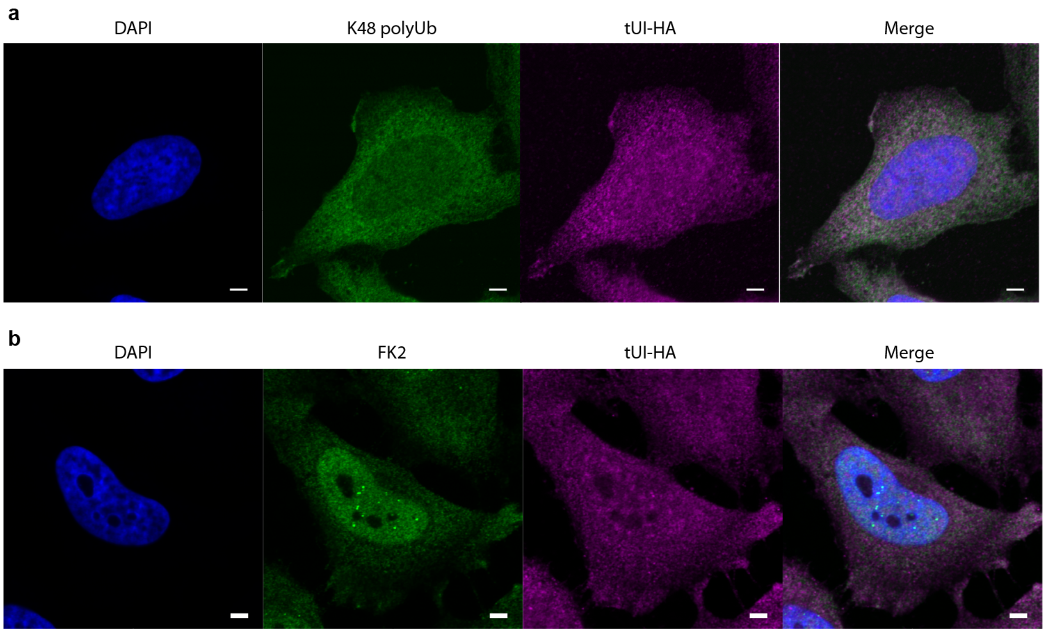


**Fig. S9. Free Ub staining by HA-tUI shows a different subcellular distribution than K48-linked polyUb or total conjugated Ub.** HeLa cells were fixed with 4% PFA and stained with DAPI (blue), anti-K48 polyUb antibody (green), and HA-tUI (magenta) in panel **a**, or DAPI (blue), anti-FK2 Ub antibody (green), HA-tUI (magenta) in panel (**b**), as described in Methods.

Supplementary Figure S10

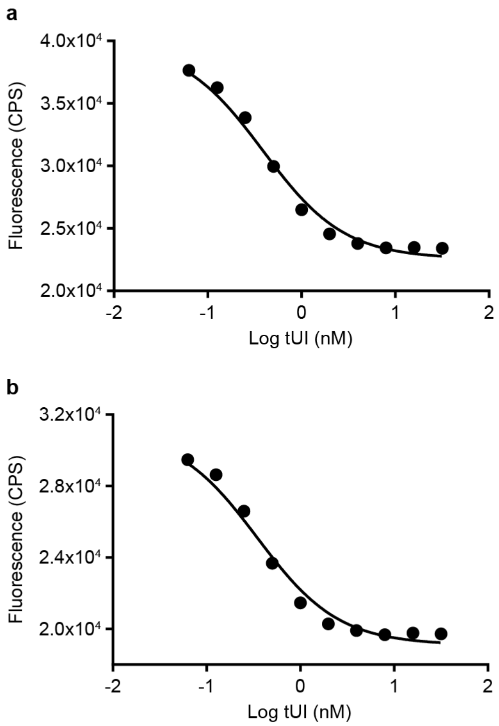


**Fig. S10. Competition binding assays to measure tUI affinity for Ub.** Competition binding assays were performed with **a**, 70 pM Atto532-tUI or **b**, 50 pM Atto532-tUI in the presence of 80 pM Ub. tUI was titrated from 0.063 nM to 32 nM, from which a $K_i$ of 194 ± 6 pM was determined.
